# PPP1R3G Deletion Blocks RIPK1-Mediated Apoptosis and Necroptosis in Doxorubicin-Induced Cardiotoxicity

**DOI:** 10.1101/2025.09.26.678784

**Authors:** Xueling Ma, Ken Chen, Zhigao Wang

## Abstract

Cardiotoxicity is a major limitation of cancer chemotherapy, exemplified by doxorubicin (DOX), yet its underlying mechanisms remain incompletely defined. Here, we identify Protein Phosphatase 1 Regulatory Subunit 3G (PPP1R3G) as a critical amplifier of DOX-induced cardiotoxicity. We show that DOX activates both apoptosis and necroptosis in vitro. Mechanistically, DOX first induces p38-dependent inhibitory phosphorylation of receptor-interacting protein kinase 1 (RIPK1), providing a transient brake on cell death. PPP1R3G facilitates the removal of inhibitory phosphorylation, thereby permitting RIPK1 activation, oligomerization, and downstream apoptotic signaling. Activated RIPK1 further promotes mitochondrial DNA (mtDNA) release, which induces IFN-β-mediated ZBP1 expression and establishes a positive feedback loop that amplifies late-stage necroptosis. Genetic ablation of *Ppp1r3g* suppresses both apoptosis and necroptosis in cardiomyocytes, attenuates inflammatory cytokine production, and protects mice from DOX-induced cardiac injury and mortality. These findings delineate a PPP1R3G-RIPK1 axis that converts an early protective phosphorylation checkpoint into sustained death signaling and identify PPP1R3G as a potential therapeutic target for cardioprotection.

## Introduction

Cardiotoxicity is among the most severe adverse effects of chemotherapy, particularly in adolescents^1,2^. Doxorubicin (DOX), a prototypical anthracycline, is highly effective against both solid and hematological malignancies and has markedly improved overall survival. Unfortunately, its clinical benefit is offset by the risk of cardiac injury^3^. The dose-dependent cardiotoxicity of DOX was first recognized more than half a century ago^4^, yet the underlying mechanisms remain incompletely understood. Therefore, advancing research is essential not only to deepen our understanding of its mechanisms, but also to identify novel therapeutic targets that ultimately improve patient outcomes.

The loss of terminally differentiated cardiomyocytes is the direct cause of cardiac dysfunction. Apoptosis is the most widely reported and studied form of cell death in DOX-induced cardiotoxicity^5-7^, which has been considered an immunologically silent process. However, recent studies have revealed that cardiac inflammation has emerged as a key pathophysiological link across the broadening spectrum of cancer therapy-related cardiotoxicities, serving as a mechanistic driver of myocarditis, cytokine release-induced cardiac dysfunction, and inflammation-associated arrhythmias^8^. Notably, pro-inflammatory cytokines such as tumor necrosis factor (TNF)-α and interleukin (IL)-6 are upregulated after DOX treatment^9^, driving sterile inflammation within damaged tissue. In contrast to apoptosis, necroptosis represents a distinct cell death pathway characterized by plasma membrane rupture, release of damage-associated molecular patterns (DAMPs), and robust immune activation, implicated in a wide range of pathological conditions^10,11^. When caspase-8 activity is inhibited or insufficient, necroptosis is preferentially activated instead of apoptosis^12,13^, and this shift may further amplify inflammation and exacerbate cardiac injury.

Receptor-interacting protein kinase 1 (RIPK1) is a central regulator of both apoptosis and necroptosis, and serves as a key modulator of inflammation^14,15^. Its kinase activity is tightly regulated through multiple mechanisms. Following TNF stimulation, p38/MK2 induces global RIPK1 phosphorylation, with several inhibitory sites constraining its kinase activity and thereby preventing cell death^16,17^. Among these, serine 25 (Ser25) is a well-characterized inhibitory site, whose phosphorylation alters the activation loop interaction^18^. In addition, phosphorylation at serine 320 (S320) in human RIPK1 (corresponding to S321 in mouse) also suppresses its activation^16,17,19^. Together, these inhibitory phosphorylation events act as essential safeguards that repress RIPK1 activity and preserve cellular homeostasis under stress and inflammatory conditions.

Through a CRISPR-based whole-genome knockout screen, our laboratory identified that Protein Phosphatase 1 Regulatory Subunit 3G (PPP1R3G), a regulatory subunit of protein phosphatase 1 (PP1), is required for RIPK1-dependent apoptosis and necroptosis. Mechanistically, PPP1R3G interacts with PP1γ to form a functional holoenzyme that dephosphorylates the inhibitory sites on RIPK1. Importantly, *Ppp1r3g*^*-/-*^ mice exhibit strong resistance to TNFα-induced systemic inflammatory response syndrome (SIRS), confirming its critical role in vivo^20^. Given the importance of RIPK1 inhibitory phosphorylations in restraining cell death, these findings suggest that PPP1R3G may serve as a key regulator of DOX-induced cardiotoxicity. In this study, we investigated the role of PPP1R3G in DOX-triggered apoptosis and necroptosis and further evaluated the cardioprotective effects of PPP1R3G deficiency in a mouse model of DOX-induced cardiotoxicity.

## Methods and materials

### Reagents

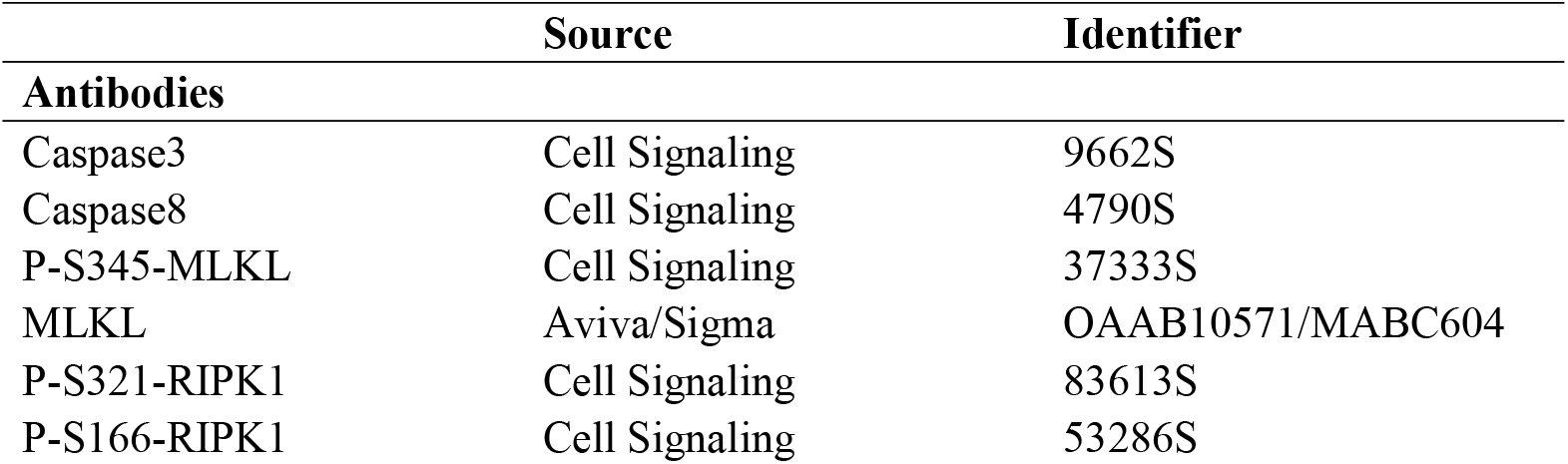
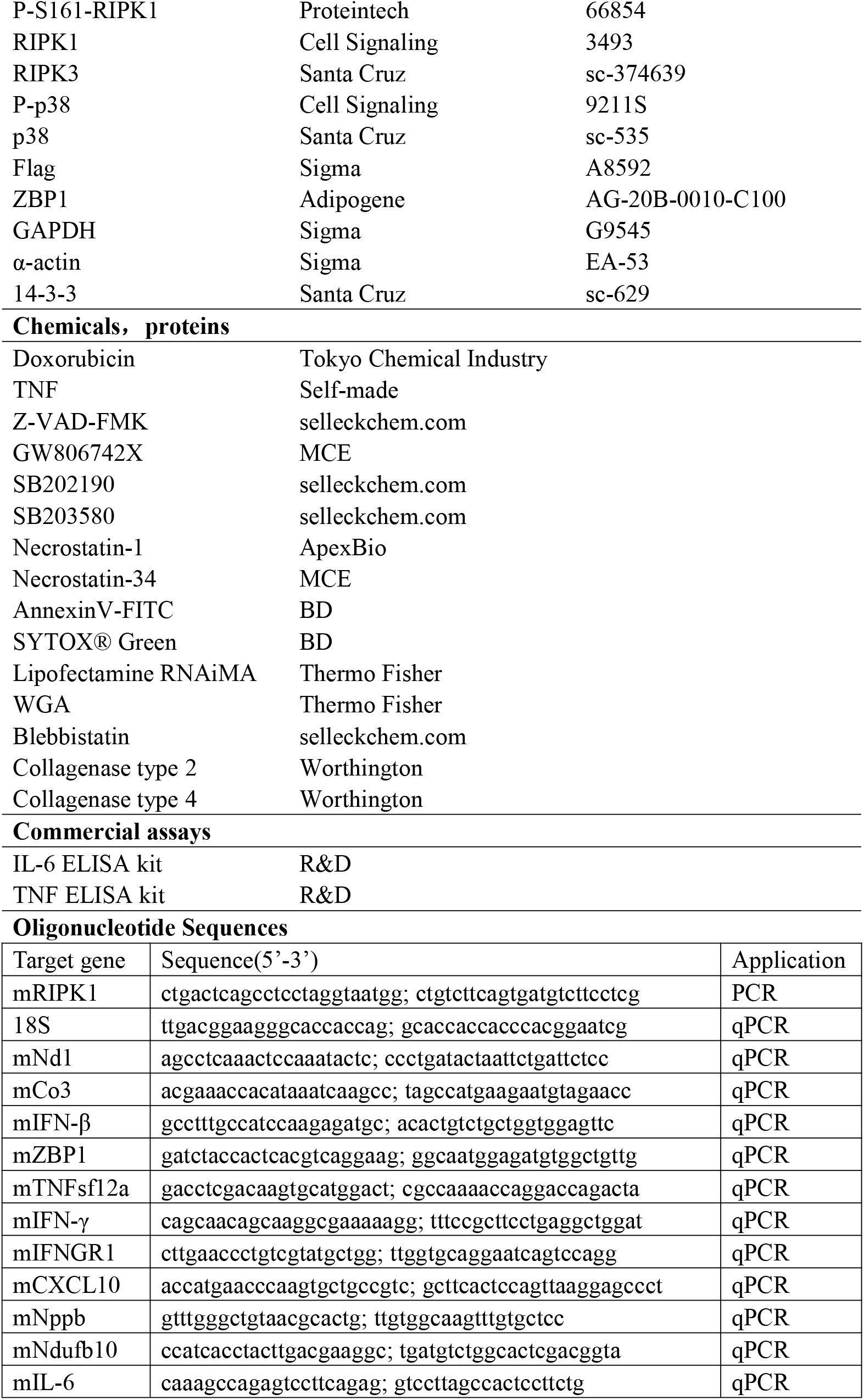
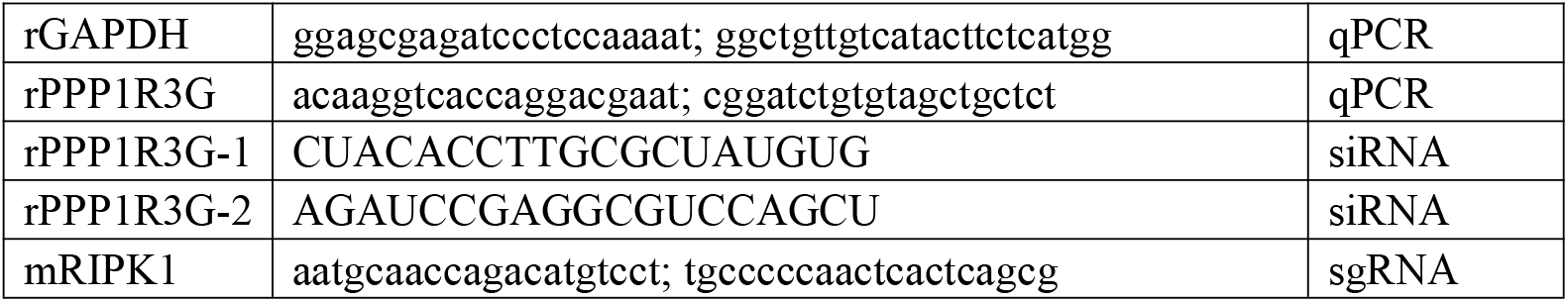

### Cell culture

H9C2 rat cardiomyoblasts (H9C2) and immortalized mouse embryonic fibroblasts (MEFs) were purchased from ATCC, and the Lenti-X™ 293T cell line was obtained from Takara. Primary wild-type and *Ppp1r3g*^*−/−*^ MEFs were isolated from E12.5-E14.5 embryos of wild-type and knockout mice following standard protocols^20^. Immortalized ZBP1-knockout (ZBP1-KO) and ZBP1-rescue MEF lines were kindly provided by the Balachandran laboratory at Fox Chase Cancer Center^21^. All cell lines were maintained in DMEM supplemented with 10% FBS, penicillin (100 U/ml), and streptomycin (100 μg/ml). Immortalized KO and rescue lines were cultured in selective medium to ensure transgene stability.

Cardiomyocyte isolation was performed as previously described^22,23^, except that 2,3-butanedione monoxime (BDM) was replaced with 5 mM blebbistatin. All cultures were maintained under standard conditions (37 °C, 5% CO_2_).

### Generation of stable cell lines

CRISPR-Cas9-mediated gene editing was used to generate RIPK1 KO or kinase domain deletion (KDD) clones in immortalized MEFs. Single-guide RNAs (sgRNAs) targeting RIPK1 were cloned into lentiviral vectors and co-transduced with Cas9 as described before^20^. Stable KO and KDD clones were selected and validated by immunoblotting and genomic DNA sequencing.

### Cell death assays

Cell viability was measured with the CellTiter-Glo Luminescent Cell Viability Assay (Promega) according to the manufacturer’s instructions. For microscopic analysis, cells were stained either with SYTOX Green and DAPI, or with Annexin V-FITC/PI and imaged using a fluorescence microscope as described before^24^.

### Western blotting

Cells were lysed in NP-40 buffer (50 mM Tris, pH 7.4, 137 mM NaCl, 1% NP-40, and 10% glycerol) containing protease and phosphatase inhibitors. After centrifugation at 15,000 rpm for 10 min, the supernatants (NP-40-soluble fractions) were collected and boiled in SDS sample buffer. The pellets (NP-40-insoluble fractions) were washed twice with PBS, then resuspended and boiled in SDS sample buffer. Equal amounts of protein were separated by SDS-PAGE and transferred to PVDF membranes, which were incubated with the indicated primary antibodies followed by HRP-conjugated secondary antibodies. Protein bands were visualized using enhanced chemiluminescence (ECL) reagents and imaged with a ChemiDoc system (Bio-Rad).

### RNA extraction and quantitative PCR

Total RNA was extracted from mouse heart tissues and cultured cells using TRIzol reagent (Invitrogen). For siRNA experiments, H9C2 cells were transfected with *Ppp1r3g* or control siRNA, and RNA was collected 48 hours post-transfection for RT-qPCR. For mouse hearts, apical tissue was rinsed in PBS, minced, and homogenized with a tissue blender prior to RNA extraction. Relative expression levels were calculated using the 2^-ΔΔCt method with 18S rRNA as the internal control.

### Cytosolic mtDNA detection by (q)PCR

Cells were fractionated at 4 °C by differential centrifugation in Buffer A (20 mM HEPES, pH 7.4, 10 mM KCl, 1.5 mM MgCl_2_, 1 mM DTT) with 250 mM sucrose. Homogenates were cleared at 500 g, 10 min (nuclear pellet), then spun at 20,000 g for 10 min to obtain the organelle pellet, then the supernatant was ultracentrifuged at 100,000 g for 1 hour to yield the cytosolic fraction. For qPCR, the cytosolic signal was normalized to the corresponding organelle fraction mtDNA amount. Endpoint PCR was performed in parallel using fraction inputs pre-normalized to equal protein concentrations to control loading; PCR products were analyzed by agarose gel electrophoresis.

### Echocardiography

Transthoracic echocardiography was performed using the Vevo 3100 high-resolution imaging system (VisualSonics, Toronto, Canada) equipped with a high-frequency small-animal transducer (MX550D, 40 MHz). Mice were anesthetized with 1-2% isoflurane and positioned supine on a heated platform with ECG and respiratory monitoring. Parasternal short-axis views were obtained, and M-mode tracings were recorded at the papillary muscle level. Left ventricular (LV) end-diastolic diameter, LV end-systolic diameter, ejection fraction (EF), fractional shortening (FS), stroke volume, and cardiac output were calculated using VevoLab software. Each parameter was averaged over two consecutive cardiac cycles, and all analyses were performed in a blinded manner.

### Histology and immunofluorescence

Mouse hearts were harvested, rinsed in PBS, and fixed in 4% paraformaldehyde at 4 °C overnight. Tissues were dehydrated, embedded in paraffin, and sectioned at 5 µm. For H&E staining, sections were deparaffinized, rehydrated, and stained with hematoxylin and eosin (AML Labs), then evaluated independently by two blinded observers. For immunofluorescence, sections underwent antigen retrieval, permeabilization, and serum blocking, followed by incubation with primary antibodies overnight at 4°C and fluorophore-conjugated secondary antibodies at room temperature. Nuclei were counterstained with DAPI. Images were acquired by fluorescence microscopy, and ≥5 fields per section were quantified using ImageJ.

### Mouse models and experimental design

*Ppp1r3g*^-/-^ mice were generated using CRISPR-Cas9-mediated deletion^20^. Male C57BL/6 wild-type (WT) and *Ppp1r3g*^*-/-*^ mice (8∼10 weeks old) were used. Doxorubicin (20 mg/kg, i.p.) was prepared in sterile water and diluted in PBS immediately before injection; control mice received PBS. Baseline echocardiography was performed 1 day before treatment, followed by imaging and tissue collection on day 6. For survival analysis, mice received DOX and were monitored without further intervention. All animal procedures were approved by the Institutional Animal Care and Use Committee (IACUC).

### Public Database Analysis

Transcriptomic datasets of doxorubicin (DOX)-treated mouse hearts (GSE226116, GSE223698) and iPSC-derived cardiomyocytes (GSE217421) were obtained from the GEO database (https://www.ncbi.nlm.nih.gov/geo/). Raw mouse RNA-seq reads were aligned to the GRCm38 reference genome using STAR (v2.7.11b) with Ensembl release 99 annotation, and gene-level counts were derived with StringTie (v2.2.3). For the iPSC-CM dataset, processed expression matrices were directly retrieved. Differential expression and gene set enrichment analyses (GSEA) were performed using the online platform (https://www.bioinformatics.com.cn)^25^, and results were visualized as volcano plots, heatmaps, and enrichment plots.

### Statistical analysis

Data are presented as mean ± SEM unless stated otherwise. Pairwise comparisons were performed using two-tailed Student’s T-tests. For experiments with a single factor and ≥3 treatment groups, one-way ANOVA was used, followed by Tukey’s post hoc test for multiple comparisons. For experiments involving two factors (e.g., genotype × treatment), two-way ANOVA with Šidák’s correction was applied. Survival data were analyzed by Kaplan–Meier curves and compared using the log-rank (Mantel–Cox) test. Statistical significance was defined as p < 0.05. Analyses were conducted in GraphPad Prism. All statistical analyses were performed using GraphPad Prism (version 10; GraphPad Software, San Diego, CA, USA).

## Results

### 1. Doxorubicin Activates TNF Signaling, Apoptotic, and Necroptotic Pathways in the Heart

We first analyzed two publicly available transcriptomic datasets of doxorubicin (DOX)-treated mice, both designed to capture early cardiotoxic responses. The GSE226116 dataset included heart tissue collected after the second DOX dose (day 1, 5 mg/kg; day 8, 5 mg/kg), whereas GSE223698 contained cardiomyocytes isolated two days after a single 25 mg/kg DOX injection. GSEA revealed significant upregulation of the TNF signaling pathway in both datasets (GSE226116: NES = 1.41, *P* = 0.003, *q* = 0.002; GSE223698: NES = 1.73, *P* = 0.003, *q* = 0.002). The cytokine-cytokine receptor interaction pathway was also consistently enriched (GSE226116: NES = 1.50, *P* < 0.001, *q* < 0.001; GSE223698: NES = 2.01, *P* < 0.001, *q* < 0.001). In addition, apoptosis was significantly enriched in GSE226116 (NES = 1.31, *P* = 0.024, *q* = 0.030), while necroptosis was significantly enriched in GSE223698 (NES = 1.54, *P* = 0.026, *q* = 0.019) (**Fig. 1A, B**). These findings suggest that DOX-induced cardiotoxicity involves activation of inflammatory, apoptotic, and necroptotic pathways. Analysis of a third dataset of iPSC-derived cardiomyocytes (GSE217421) further supported this conclusion, showing enrichment of cytokine-cytokine receptor interaction, necroptosis, and TGF-β signaling pathways, suggesting similar responses to DOX-treatment in human cardiomyocytes (**Fig. S1A**). Building on these transcriptomic insights, we next validated the involvement of apoptosis and necroptosis in DOX-induced cardiotoxicity using H9C2 cardiomyoblasts. DOX treatment reduced H9C2 cell viability in a dose-dependent manner, with approximately 30% cell death observed after 16 h exposure to 60 μM DOX (**Fig. 1C**). Co-treatment with TNF, a key initiator of cell death, further exacerbated DOX-induced cytotoxicity (**Fig. 1D**). Western blotting showed cleavage of caspase-8 and caspase-3, confirming the activation of apoptosis (**Fig. 1E**). Inhibition of apoptosis by Z-VAD effectively blocked caspase activation (**Fig. 1E**) but paradoxically increased overall cell death (**Fig. 1D**). To explore the underlying mechanisms, cells were treated with various inhibitors. NAC (N-acetylcysteine), an antioxidant, attenuated cell death induced by DOX or DOX + TNF, confirming the role of reactive oxygen species (ROS) in activating DOX-induced cell death^6^. Importantly, the MLKL inhibitors GW806742 and Saracatinib markedly reduced death in the DOX + TNF + Z-VAD group (**Fig. 1F**), without affecting caspase-8 or caspase-3 cleavage (**Fig. 1G**), suggesting a shift toward necroptosis when apoptosis was blocked. GW806742 provided protection even at very low concentrations (**Fig. S1D**). Annexin V/PI staining further confirmed its protective effect on both apoptotic and total dead cells, with incomplete overlap between Annexin V and PI signals (**Fig. 1H-J**). Together, these results suggest that DOX may trigger cardiomyocyte death directly, as well as exacerbate injury by enhancing cytokine-mediated inflammatory signaling (**Fig. 1K)**.

**Fig 1.**
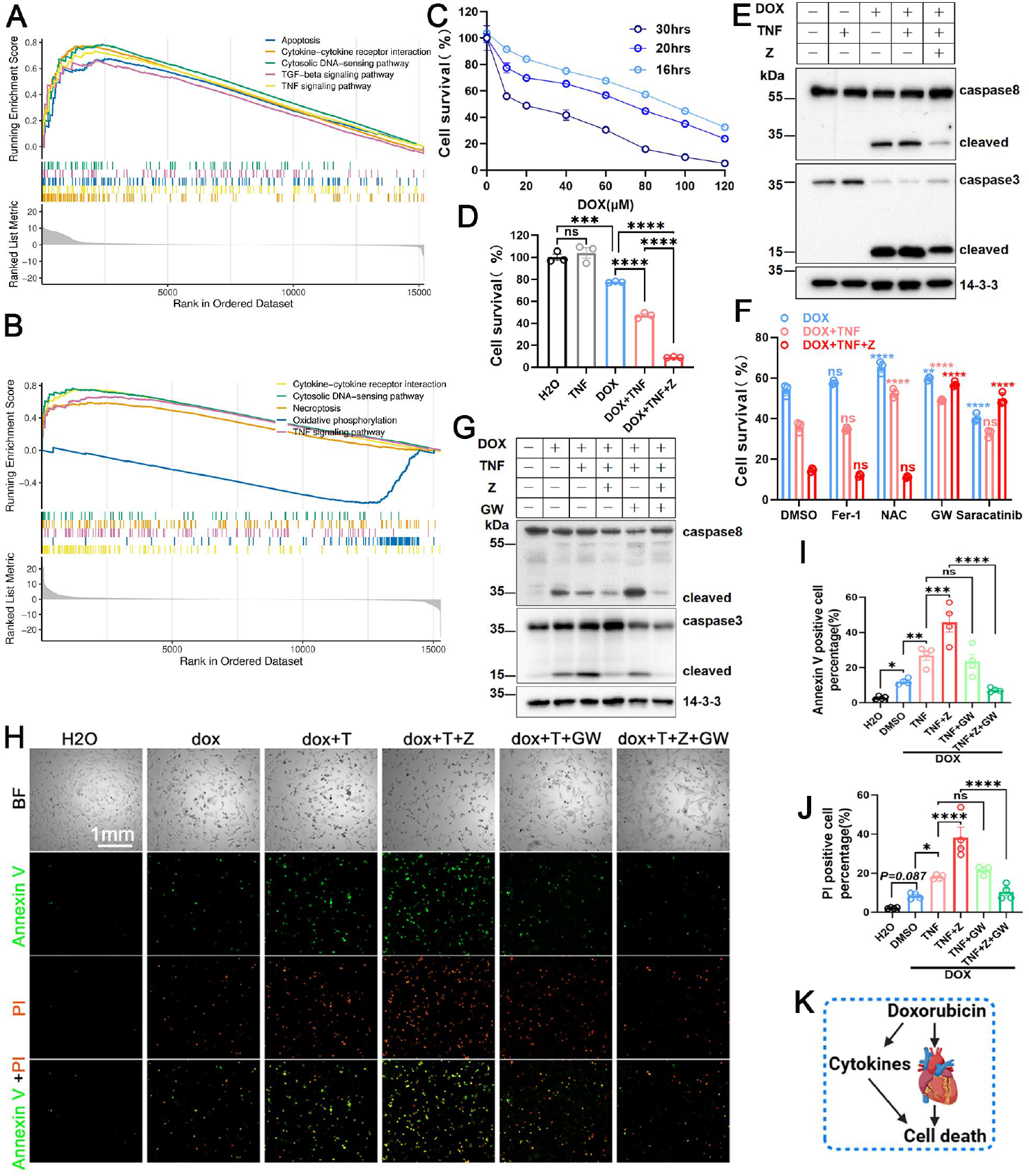
Doxorubicin Activates TNF Signaling, Apoptotic, and Necroptotic Pathways in the Heart. **(A, B)** GSEA of two transcriptomic datasets. Analysis of GSE226116 (mouse hearts) and GSE223698 (primary cardiomyocytes) was performed using the mouse KEGG gene sets. Normalized enrichment score (NES), nominal P value, and FDR q value are reported in the Results. **(C)** Cell survival assessed by CellTiter-Glo assay of H9C2 treated with different doses of DOX for different durations. **(D)** H9C2 cells were exposed to DOX (60 μM, 16 hours), and cell survival was assessed by CellTiter-Glo assay. Z-VAD was used at 20 μM. TNF was applied at 6 ng/mL when used alone, but reduced to 2 ng/mL in combination with Z-VAD to avoid excessive cell death; the same concentrations were applied throughout the study. **(E)** Immunoblot analysis of apoptotic markers in H9C2 cells treated with DOX ± TNF ± Z (60 μM DOX, 7 hours). **(F)** Viability of H9C2 cells treated with DOX ± TNF ± Z in the presence of NAC (2 mM), GW806742 (10 μM), or Saracatinib (5 μM), assessed by CellTiter-Glo assay. **(G)** Immunoblot analysis of apoptotic markers in H9C2 cells treated with DOX ± TNF ± Z with or without GW806742. **(H)** Representative images of Annexin V/PI staining in H9C2 cells treated for 12 hours. A shorter incubation time was chosen for live-cell staining compared with CellTiter-Glo assays to prevent loss of dead cells. **(I, J)** Quantification of Annexin V/PI staining shown in **(H)** (n = 3 technical replicates per condition). **(K)** Working model. DOX may trigger cardiomyocyte death directly, or indirectly through cytokine-mediated inflammatory signaling.

### 2. Loss of PPP1R3G alleviates doxorubicin-induced cell death

Our previous studies identified PPP1R3G as a critical regulator of both apoptosis and necroptosis^20^. We therefore investigated whether loss of PPP1R3G could attenuate DOX-induced cell death. Silencing PPP1R3G with siRNA in H9C2 cells significantly reduced cell death induced by DOX/DOX+TNF/DOX+TNF+Z, with the second siRNA construct achieving more efficient knockdown and stronger protective effects **(Fig. 2A)**. Consistently, *Ppp1r3g*^*-/-*^ MEFs exhibited resistance to DOX-induced cell death, with or without TNF (**Fig. 2B-D, Fig. S2A**). Western blot analysis confirmed that *Ppp1r3g* deficiency suppressed both necroptosis and apoptosis, as evidenced by reduced p-S345-MLKL, decreased caspase-3 cleavage, and less accumulation of insoluble MLKL and RIPK3 (**Fig. 2E, F**). Importantly, primary cardiomyocytes isolated from adult *Ppp1r3g*^*-/-*^ mice were also protected against DOX-induced cell death, showing a markedly lower proportion of Sytox Green-positive cells compared with wild-type controls (**Fig. 2G, H**). Overall, *Ppp1r3g* knockout reduces DOX-induced apoptosis and necroptosis in H9C2 cells, MEFs, and notably in cardiomyocytes (**Fig. 2I**).

**Fig 2.**
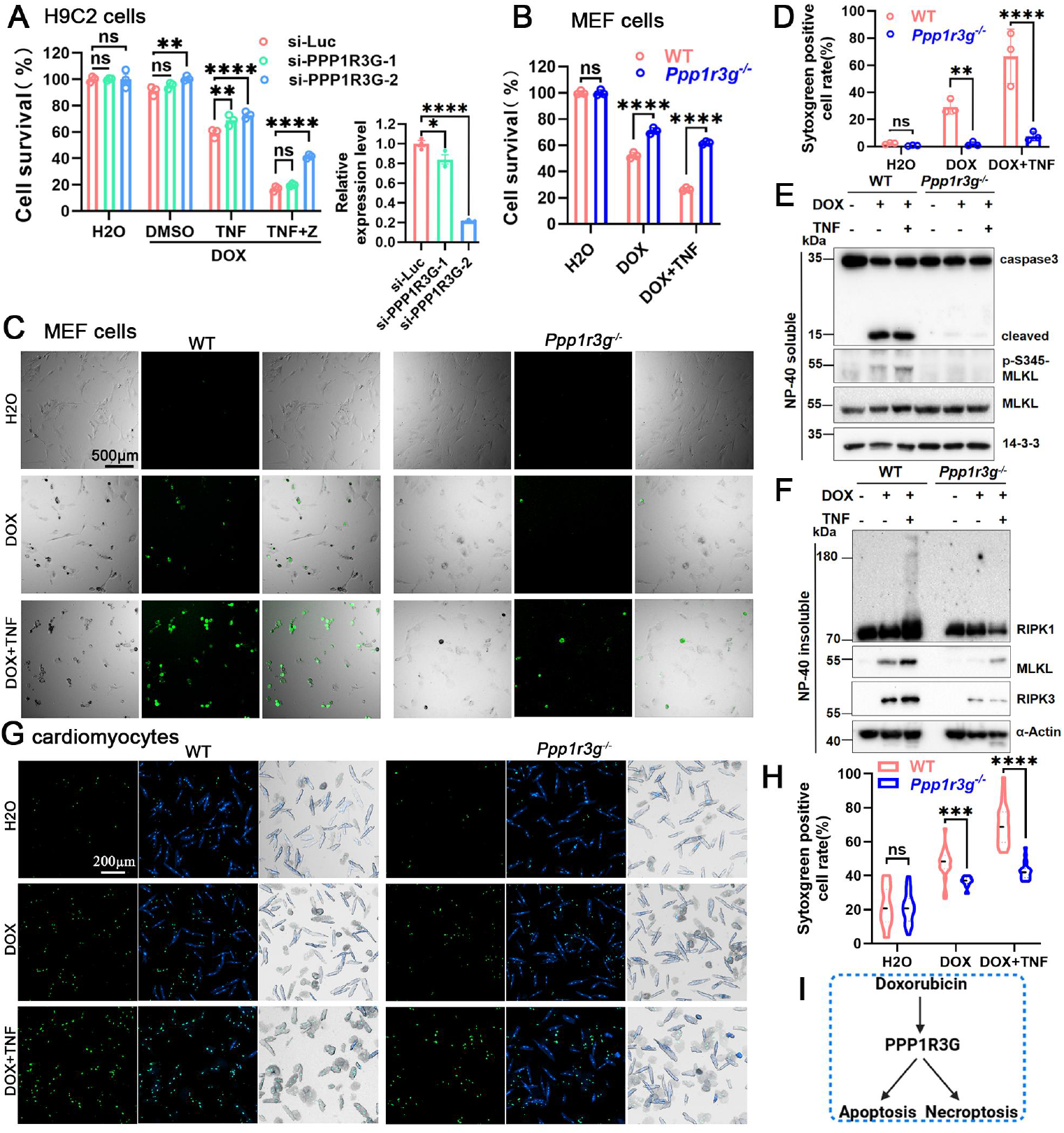
Loss of PPP1R3G alleviates doxorubicin-induced cell death. **(A)** PPP1R3G knockdown in H9C2 cells attenuated DOX-induced cell death. Cell viability was measured by CellTiter-Glo assay after siRNA-mediated PPP1R3G silencing and DOX ± TNF ± Z treatment for 36 hours; knockdown efficiency is shown in the right panel. **(B)** Cell survival of WT and *Ppp1r3g*^*−/−*^ MEFs. Cells were treated with DOX (30 μM) ± TNF for 24 hours and analyzed by CellTiter-Glo assay. The same concentration of DOX was used for all MEF experiments. **(C)** Representative images of Sytox Green staining of WT and *Ppp1r3g*^*−/−*^ MEFs treated with DOX ± TNF for 12 hours. **(D)** Quantification of Sytox Green-positive cells shown in **(C). (E, F)** Immunoblot of apoptotic and necroptotic markers in NP-40 soluble and insoluble fractions in WT and *Ppp1r3g*^*−/−*^ MEFs. MEFs were treated with DOX ± TNF for 12 hours. **(G)** Representative images of Sytox Green staining of WT and *Ppp1r3g*^*−/−*^ primary cardiomyocytes treated with DOX ± TNF. **(H)** Quantification of Sytox Green-positive cells shown in **(G). (I)** Working model. *Ppp1r3g* knockout reduces DOX-induced apoptosis and necroptosis.

### 3. PPP1R3G promotes DOX-induced apoptosis and necroptosis by regulating RIPK1 phosphorylation, activation, and oligomerization

Given our previous findings that PPP1R3G regulates RIPK1 kinase activity and thereby regulating apoptosis and necroptosis^20^, we next investigated whether PPP1R3G regulates DOX-induced cell death through modulation of RIPK1 kinase activity. Treatment of H9C2 cells with the RIPK1 inhibitor necrostatin-1 (Nec-1) attenuated DOX-induced cell death (**Fig. S3A**). Western blotting revealed that DOX plus TNF triggered RIPK1 activation, indicated by p-S166-RIPK1, and Z-VAD further enhanced this activation (**Fig. S3B**). Notably, in both wild-type and *Ppp1r3g*^*-/-*^ MEFs, another auto-activating site S161 of RIPK1 was phosphorylated. Interestingly, p-S161-RIPK1 is reported to be activated by mitochondrial ROS^26^, which increase following DOX treatment^6^. Importantly, PPP1R3G deficiency maintained higher levels of inhibitory phosphorylation of RIPK1 (p-S321), accompanied by a corresponding reduction in RIPK1 activation, as indicated by lower p-S161 (**Fig. 3A**). Since inhibitory phosphorylation of RIPK1 is regulated by p38^16,17^, we pretreated cells with two p38 inhibitors and found that inhibition of p38 increased DOX-induced cell death (**Fig. 3B**), indicating that inhibitory phosphorylation of RIPK1 is critical for restraining DOX-triggered cytotoxicity. Time-course analysis further revealed that DOX treatment induced p38 phosphorylation, followed by inhibitory phosphorylation of RIPK1 (p-S321), representing an early cellular attempt to limit cell death. However, as treatment progressed, inhibitory phosphorylation declined, RIPK1 activation increased, and both apoptosis (cleaved caspase-3) and necroptosis (p-MLKL) were triggered (**Fig. 3C**).

**Fig 3.**
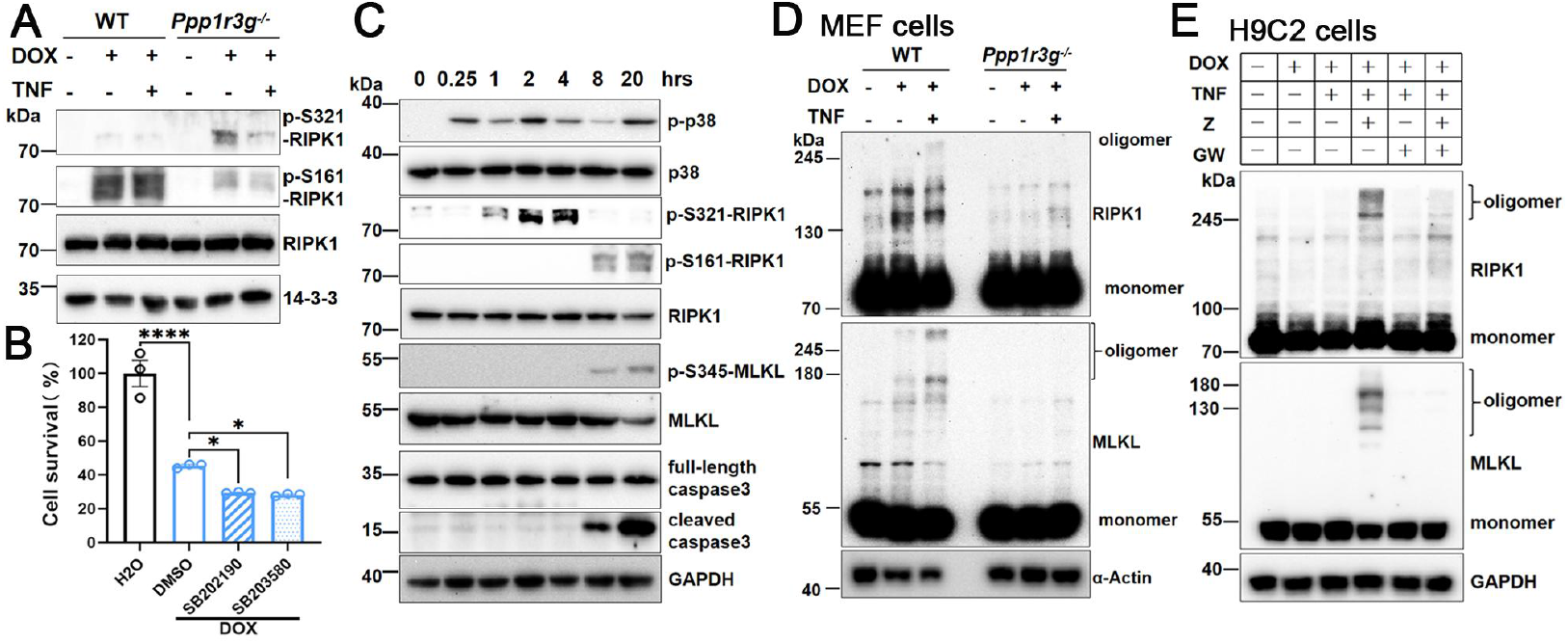
PPP1R3G promotes DOX-induced apoptosis and necroptosis by regulating RIPK1 phosphorylation, activation, and oligomerization. **(A)** Immunoblotting of WT and *Ppp1r3g*^*−/−*^ MEFs treated with DOX ± TNF for 16 hours. **(B)** Cell survival of MEFs pretreated with p38 inhibitors (SB202190, SB203580) followed by DOX treatment, analyzed by CellTiter-Glo assay. **(C)** Time-course immunoblot analysis of RIPK1 phosphorylation in DOX-treated MEFs. **(D)** Non-reducing immunoblot analysis of RIPK1 and MLKL oligomerization in WT and ***Ppp1r3g***^***−/−***^ MEFs treated with DOX ± TNF for 16 h. **(E)** Non-reducing immunoblot analysis of RIPK1 and MLKL oligomerization in H9C2 cells treated with DOX ± TNF ± Z ± GW806742.

Since RIPK1 oligomerization is a prerequisite for downstream signaling, we performed a non-reducing gel and found that DOX+TNF+Z treatment induced RIPK1 and MLKL oligomerization in MEF cells, while PPP1R3G deficiency markedly reduced both RIPK1 and MLKL oligomerization (**Fig. 3D)**. Similarly, RIPK1 and MLKL oligomerization were observed in DOX-treated H9C2 cells, which was diminished by GW806742 treatment (**Fig. 3E**).

### 4. RIPK1 kinase activity is essential for DOX-induced Cell death

Given the critical role of RIPK1 in DOX-induced cell death, we attempted to generate RIPK1 knockout cells. As shown in **Fig.4A**, CRISPR/Cas9 was employed with two sgRNAs designed to target the 5’ coding sequence of *Ripk1*, thereby introducing frameshift mutations that disrupt protein expression. Unexpectedly, many resulting colonies did not completely lose RIPK1 expression but instead produced truncated forms of RIPK1(**Fig.4B, S4B**). Phenotypic characterization of these colonies revealed that all these colonies were resistant to T/S/Z-induced cell death, colonies with complete RIPK1 deletion (*Ripk1*^*-/-*^) exhibited increased sensitivity to DOX compared with WT cells, whereas colonies harboring a kinase-domain deletion (*Ripk1*^*KDD*^) were more resistant to DOX-induced death. (**Fig. 4C**). RIPK1 overexpression in 293T cells triggers auto-phosphorylation of S166, whereas the kinase-domain deletion construct failed to do so, suggesting loss of kinase activity (**Fig. 4D**). We speculate that RIPK1-KDD contributes to pro-survival signaling through its scaffold function, similar to previous reported other kinase-dead mutants, such as K45M, K45A, and D138N^27-29^. Indeed, *Ripk1*^*KDD*^ cells exhibited a marked reduction in caspase-3 cleavage, p-S345-MLKL, and the formation of NP-40-insoluble MLKL/RIPK3 aggregates, confirming reduction of cell death. (**Fig. 4E**).

**Fig 4.**
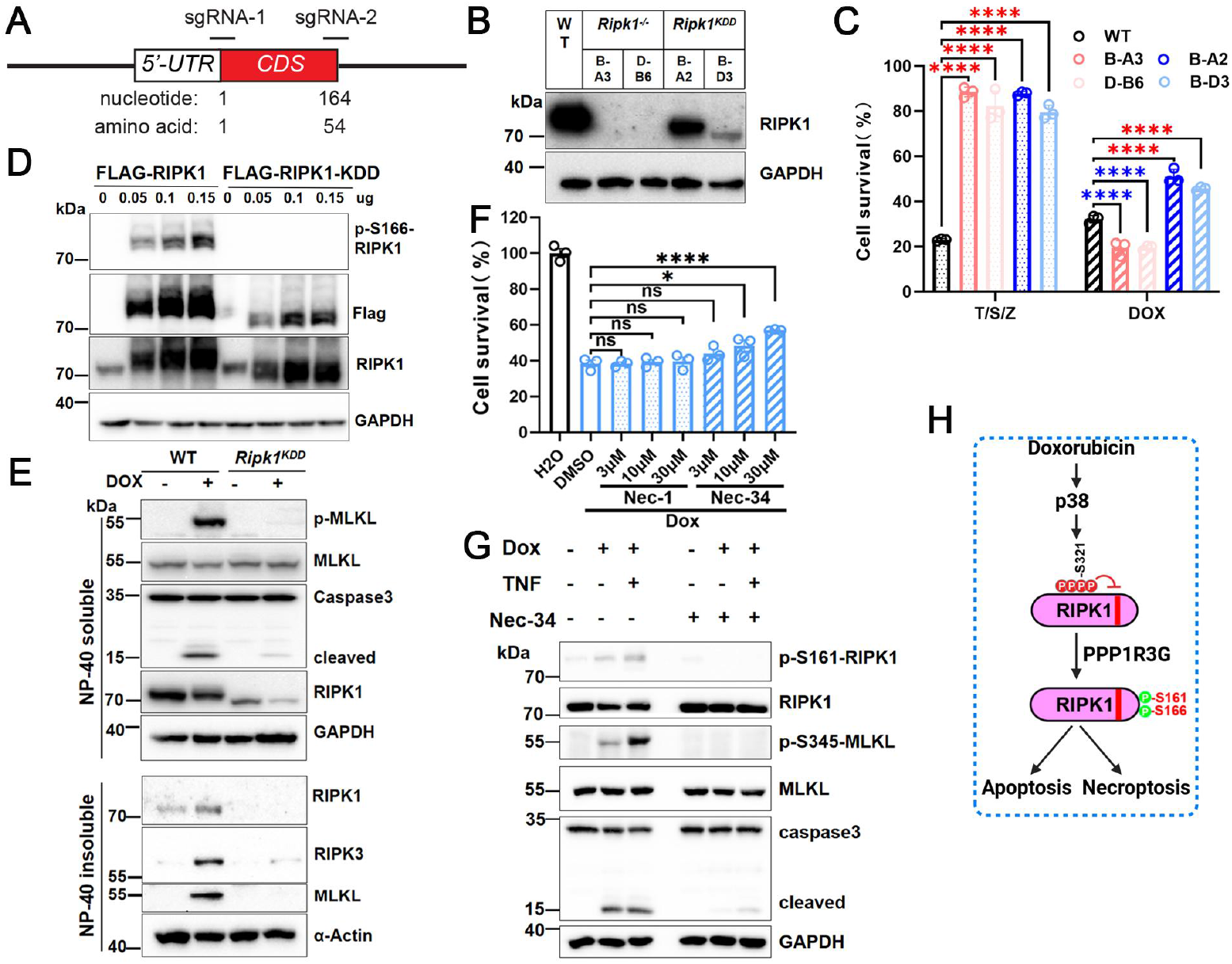
RIPK1 kinase activity is essential for DOX-induced Cell death. **(A)** Strategy for CRISPR/Cas9-mediated knockout of mouse *Ripk1*. **(B)** RIPK1 expression in individual MEF colonies generated by CRISPR-mediated *Ripk1* knockout. **(C)** Cell viability of MEF colonies treated with T/S/Z or DOX, assessed by CellTiter-Glo assay. **(D)** Immunoblot of 293T cells transfected with increasing doses of FLAG-RIPK1(WT) or FLAF-RIPK1-KDDplasmids. **(E)** Immunoblot analysis of DOX-treated MEFs.The *Ripk1*^*-/-*^ clone B-A3 and *Ripk1*^*KDD*^ clone B-A3 were used in all subsequent experiments. **(F)** Cell viability of WT MEFs treated with DOX ± Nec-1 or Nec-34, analyzed by CellTiter-Glo assay. **(G)** Immunoblot of WT MEFs treated with DOX ± Nec-34. **(H)** Working model. DOX activates p38 to induce inhibitory phosphorylation of RIPK1, which is later removed by PPP1R3G, permitting RIPK1 activation, oligomerization, and the execution of apoptosis and necroptosis.

To further assess the pharmacological inhibition of RIPK1 activation, we tested whether Nec-1 could block DOX-induced cell death in MEFs. Surprisingly, Nec-1 showed no protective effect, whereas necrostatin-34 (Nec-34) effectively suppressed DOX-induced cell death (**Fig. 4F**). The limited efficacy of Nec-1 validates a previous report that the phosphorylation of S161 alters the activation loop’s conformation, which blocks Nec-1 binding but doesn’t affect Nec-34 binding^30^. Furthermore, Western blotting confirmed that Nec-34 attenuated p-S161 activation and decreased both apoptosis and necroptosis (**Fig. 4G**).

Together, these findings support a model in which DOX induces ROS production, which activates p38, leading to inhibitory phosphorylation of RIPK1. This inhibitory phosphorylation is subsequently removed by PPP1R3G, thereby enabling RIPK1 activation, oligomerization, and induction of apoptosis and necroptosis in response to DOX (**Fig. 4H**).

### 5. RIPK1 activation promotes mitochondrial DNA release, thereby inducing ZBP1 expression and contributing to late-stage DOX-induced necroptosis

As shown in **Fig. 1A** and **Fig. 1B**, the cytosolic DNA-sensing pathway was consistently activated in both isolated cardiomyocytes (NES = 1.88, *P* < 0.001, *q* < 0.001) and heart tissue (NES = 1.50, *P* = 0.001, *q* = 0.001). Further analysis of the GSE223698 dataset revealed significant upregulation of ZBP1 transcripts in DOX-treated cardiomyocytes **(Fig. 5A, S5A)**. The core enrichment genes of the cytosolic DNA-sensing pathway in both datasets are listed in **Supplementary Table 1** and **Table 2**, and visualized in **Fig. S5B, C**. Since ZBP1 is a critical regulator of the necroptosis pathway, we next examined its role in DOX-induced cell death by comparing ZBP1-KO and ZBP1-rescued MEFs. Although early cell death was similar between the two groups, ZBP1 deficiency significantly reduced cell death at 24-36 h, indicating a late contribution of ZBP1 (**Fig. 5B**). Time-course experiments revealed that DOX induced caspase-3 cleavage as early as 6 hours. In contrast, strong ZBP1 expression wasn’t induced until 12 hours, coinciding with the appearance of necroptosis markers, such as p-S345-MLKL and accumulation of insoluble MLKL and RIPK3. Significantly, both ZBP1 induction and necroptosis markers were abrogated in *Ppp1r3g*^*-/-*^ cells (**Fig. 5C, D**). Additional TNF stimulation further elevated ZBP1 levels (**Fig. 5E**). Importantly, western blotting demonstrated that ZBP1 deletion did not affect apoptosis but impaired necroptosis, as reflected by reduced p-S345-MLKL and diminished MLKL oligomerization, again suggesting a late contribution of ZBP1 to necroptosis induction (**Fig. 5F**).

**Fig 5.**
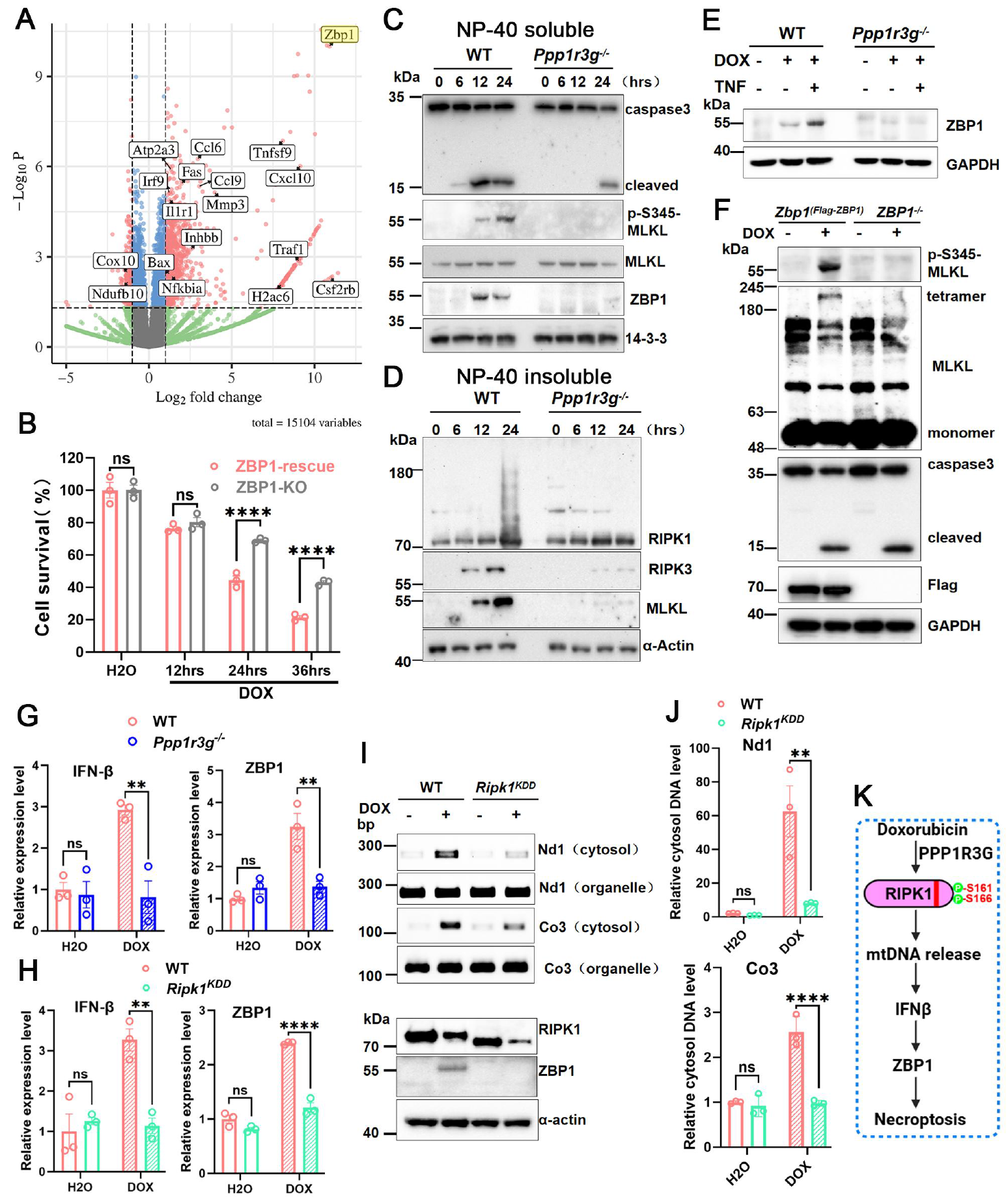
RIPK1 activation promotes mtDNA release and ZBP1-dependent late necroptosis. **(A)** Volcano plot of RNA-seq data from GSE223698. Selected differentially expressed genes associated with inflammation, cell death, and oxidative phosphorylation were labeled. **(B)** *Zbp1*^*−/−*^ and *Zbp1*^*(Flag-ZBP1)*^ MEFs treated with DOX for the indicated times and cell death was **assessed by CellTiter-Glo assay. (C, D)** Time-course immunoblot analysis of the NP-40 soluble and insoluble fractions in WT and *Ppp1r3g*^*−/−*^ MEFs treated with DOX. **(E)** Immunoblot analysis of WT and *Ppp1r3g*^*−/−*^ MEFs treated with DOX ± TNF. **(F)** Immunoblot analysis of *Zbp1*^*−/−*^ and *Zbp1*^*(Flag-ZBP1)*^ MEFs treated with DOX. **(G)** RT-qPCR analysis of IFN-β and ZBP1 mRNA levels in WT and *Ppp1r3g*^*−/−*^ MEFs treated with DOX. **(H)** RT-qPCR analysis of IFN-β and ZBP1 mRNA levels in WT and ***Ripk1***^***KDD***^MEFs treated with DOX. **(I, J)** PCR and qPCR detection of cytosolic mtDNA (Nd1 and Co3) in WT and ***Ripk1***^***KDD***^cells after DOX treatment. **(K)** Working model. RIPK1 activation promotes mtDNA release and subsequent IFN-β expression, which induces ZBP1 expression and thereby mediates late-stage DOX-induced necroptosis.

Mechanistically, DOX is known to cause mitochondrial DNA (mtDNA) release^31^, which can activate the cGAS-STING (cyclic GMP-AMP synthase-stimulator of interferon genes) pathway and promote IFN-β expression^32^. Given that ZBP1 is IFN-inducible, we assessed IFN-β and ZBP1 mRNA levels in WT, *Ppp1r3g*^*-/-*^, and *Ripk1*^*KDD*^ cells. DOX markedly induced IFN-β and ZBP1 in WT cells, but this induction was blunted by *Ppp1r3g* deficiency or RIPK1 kinase deletion, suggesting that both PPP1R3G and RIPK1 kinase activity are required for DOX-induced IFN-β and ZBP1 expression (**Fig. 5G, H**). PCR and qPCR further showed that RIPK1 kinase deletion suppressed cytosolic release of mtDNA (Nd1 and Co3) following DOX treatment (**Fig. 5I, J**). Together, these findings support a model in which PPP1R3G-dependent RIPK1 activation drives mtDNA release and IFN-β expression, leading to ZBP1 induction and thereby mediating late-stage DOX-induced necroptosis **(Fig. 5K)**.

### 6. Ppp1r3g knockout protects the mice from DOX-induced cardiotoxicity

Transcriptomic analysis of dataset GSE226116 from doxorubicin (DOX)-treated mouse hearts revealed that DOX exposure upregulated multiple inflammation-related genes, including Ccl2 and Ccl6, as well as prototypical stress-response genes such as Nppa and Nppb (**Fig. S6A**). Given that in vivo deletion of *Ppp1r3g* attenuates TNF-induced inflammation and suppresses necroptosis, we next employed an acute DOX injection model^33,34^ to investigate whether *Ppp1r3g* deficiency could mitigate DOX-induced inflammation and cardiotoxicity (**Fig. 6A**). Echocardiographic analysis six days after DOX administration showed a significant reduction in left ventricular diameter and cardiac output in WT mice (**Fig. 6B-C, S6B**). Consistently, gross pathology revealed a marked decrease in heart size in WT mice following DOX treatment, which was alleviated in *Ppp1r3g*^*-/-*^ mice (**Fig. 6D, E**). Histological analysis further demonstrated extensive vacuolization in cardiomyocytes of WT hearts by H&E staining (**Fig. S6C**), and WGA staining showed a more pronounced reduction in cardiomyocyte size in WT mice compared with knockouts (**Fig. S6D, E**). Importantly, *Ppp1r3g* deletion significantly improved survival after acute DOX exposure (**Fig. 6F**).

**Fig 6.**
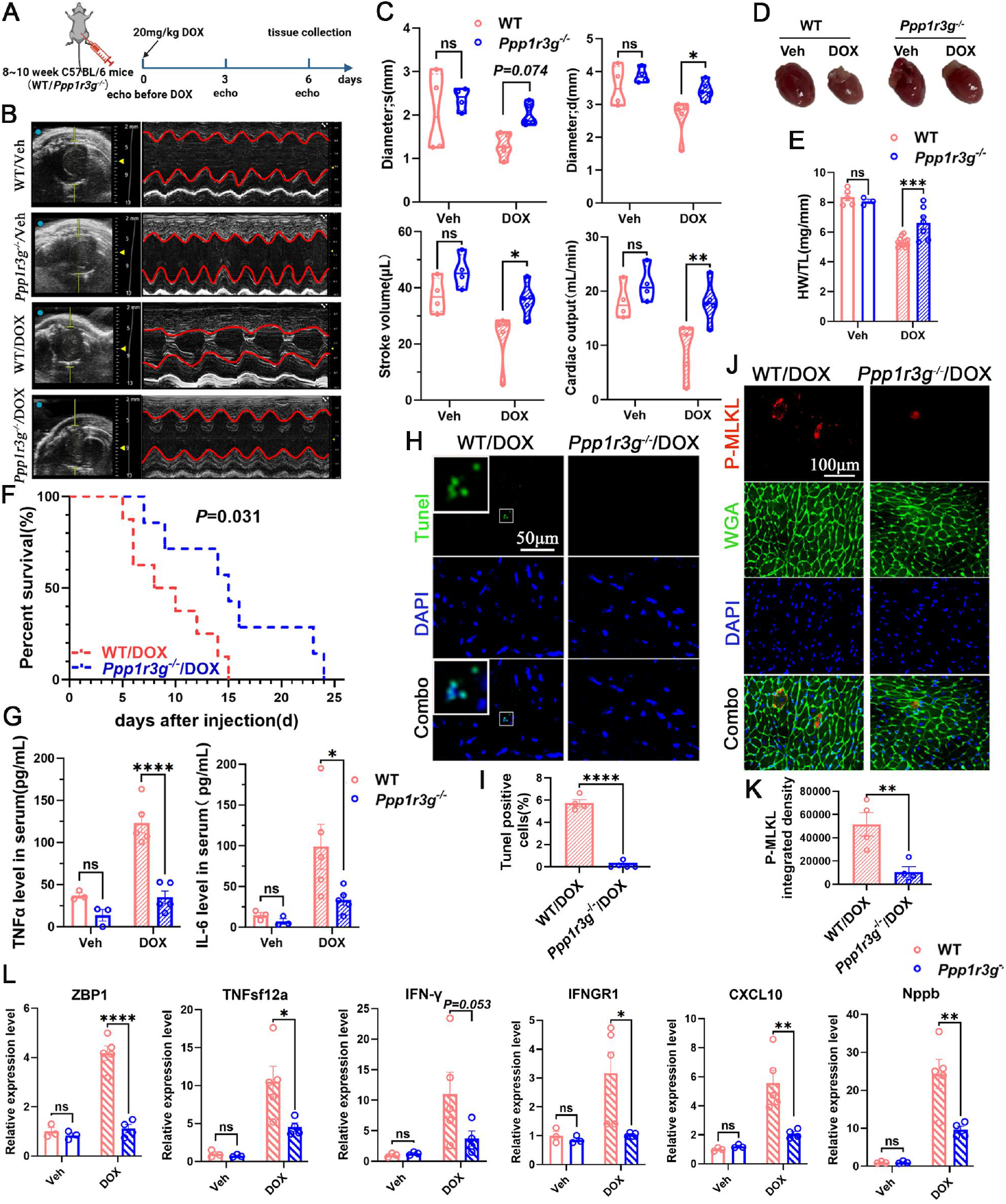
Ppp1r3g deficiency protects against DOX-induced cardiotoxicity in vivo. **(A)** Schematic of the DOX-induced acute cardiotoxicity model. **(B, C)** Echocardiographic analysis of WT and *Ppp1r3g*^*−/−*^ mice at day 6 post-DOX treatment. (**D, E**) Gross morphology of hearts and heart weight/tibia length ratio in WT and *Ppp1r3g*^*−/−*^ mice. **(F)** Kaplan-Meier survival analysis of WT and *Ppp1r3g*^*−/−*^ mice after DOX administration. (**G**) Serum TNFα and IL-6 levels in WT and *Ppp1r3g*^*−/−*^ mice following DOX treatment, assessed by ELISA. (**H, I**) TUNEL staining of cardiac sections showing apoptotic cardiomyocytes in WT and *Ppp1r3g*^*−/−*^ hearts. (**J, K**) Immunostaining for p-S345-MLKL in WT and *Ppp1r3g*^*−/−*^ hearts. (**L**) RT-qPCR analysis of apical heart tissue from WT and *Ppp1r3g*^*−/−*^ mice.

Serum cytokine analysis showed that DOX-treated WT mice exhibited a marked increase in inflammatory mediators such as TNFα and IL-6, whereas this response was blunted in *Ppp1r3g*^*-/-*^ mice (**Fig. 6G**). Given that TNF exacerbates DOX-induced cell death, this inflammatory amplification may further aggravate cardiac injury in wild-type animals. Remarkably, TUNEL staining revealed a higher proportion of apoptotic cardiomyocytes in WT mice (**Fig. 6H, I**), and p-S345-MLKL staining indicated more necroptotic cardiomyocytes in WT hearts compared with *Ppp1r3g*^*-/-*^ mice (**Fig. 6J, K**). Finally, consistent with the transcriptomic findings, RT-qPCR analysis of cardiac tissue showed that DOX treatment markedly upregulated expression of ZBP1, cytokines or cytokine receptors (TNFsf12a, IFN-γ, and IFNGR1, CXCL10), as well as the stress-response gene Nppb. These changes were significantly attenuated by *Ppp1r3g* deletion (**Fig. 6L, S6F**).

In summary, DOX exposure activates p38 signaling, leading to the global phosphorylation of RIPK1. PPP1R3G subsequently facilitates the removal of inhibitory phosphorylation from RIPK1, thereby permitting its activation (p-S161/S166) and oligomerization, which promotes downstream apoptosis at early stages. At later stages, activated RIPK1 triggers mitochondrial DNA release, leading to IFN-β production and ZBP1 upregulation, thereby inducing and amplifying necroptosis. In contrast, loss of PPP1R3G preserves RIPK1 inhibitory phosphorylation, restrains RIPK1 activation and mtDNA release, attenuates both apoptosis and necroptosis, and ultimately protects against DOX-induced cardiotoxicity in vivo **(Fig. 7)**.

**Figure 7.**
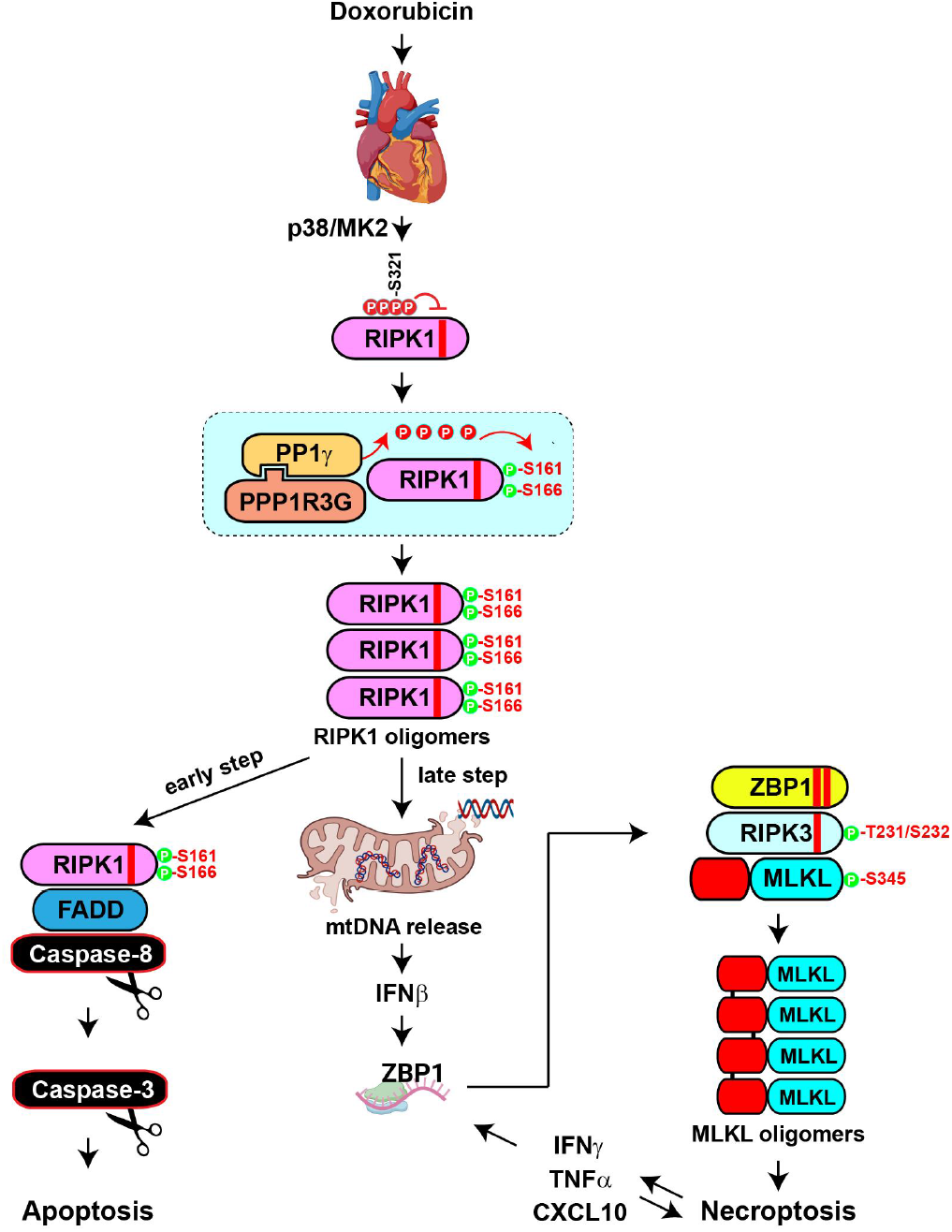
Working model. DOX promotes ROS production to activate p38 signaling which induces RIPK1 inhibitory phosphorylation.PPP1R3Gpromotes the removal of inhibitory phosphorylation of RIPK1, enabling RIPK1 activation and oligomerization. Activated RIPK1 drives early apoptosis and, at later stages, triggers mitochondrial DNA release, IFN-β production, and ZBP1 upregulation, thereby amplifying necroptosis. Loss of PPP1R3G preserves RIPK1 inhibition, limits apoptosis and necroptosis, and protects against DOX-induced cardiotoxicity.

## Disscussion

In this study, we demonstrate that DOX, a widely used chemotherapeutic, activates both apoptotic and necroptotic pathways in cardiomyocytes. We identify PPP1R3G as a central driver of DOX-induced cardiotoxicity by regulating RIPK1 activation. Mechanistically, PPP1R3G deficiency maintains inhibitory phosphorylation on RIPK1, thereby constraining its kinase activity. Because RIPK1 acts as both a kinase and a scaffold, our observation is that deletion of the RIPK1 kinase domain, but not complete RIPK1 knockout, confers protection against DOX. We further demonstrate that RIPK1 activation triggers mtDNA release, which induces ZBP1 expression and fuels a self-reinforcing necroptotic program. Importantly, in vivo deletion of *Ppp1r3g* decreases cardiomyocyte death and improves animal survival after DOX, highlighting PPP1R3G as a potential therapeutic target to mitigate anthracycline cardiotoxicity.

Apoptosis is an established contributor to DOX cardiotoxicity. DOX triggers intrinsic apoptotic pathways via oxidative stress and TOP2β-dependent DNA damage^6^, and also enhances extrinsic death-receptor signaling^35^. Interestingly, our results showed blocking apoptosis with Z-VAD paradoxically increased total cell death when TNF is present, which was rescued by MLKL inhibition, indicating that apoptosis and necroptosis function as compensatory cell death pathways in the setting of DOX-induced toxicity. Given RIPK1’s central role at the intersection of cell death pathways, precise tuning of RIPK1 activity carries therapeutic promise for diseases in which dysregulated apoptosis and/or necroptosis drive pathology^29,36^.

It is well-known that ROS are a major cause of DOX-induced cardiotoxicity. Notably, ROS also serves as a critical regulator in TNF-induced necroptotic signaling and cell death^37,38^. On the one hand, ROS scavengers such as butylated hydroxyanisole and N-acetylcysteine significantly attenuate the assembly of the RIPK1/RIPK3 necrosome and the phosphorylation of MLKL; on the other hand, silencing RIPK1 or RIPK3 can also reduce ROS production. These results together support a positive feedback loop between ROS and necroptosis^26,39^. Targeting RIPK1 as well as PPP1R3G/PP1γ thus has the potential to blunt both ROS amplification and necrosome formation in DOX-injured hearts.

A previous study reported that three crucial cysteines in RIPK1 are required for sensing ROS, and ROS subsequently activates RIPK1 autophosphorylation on serine residue 161 (S161). This specific phosphorylation then enables RIPK1 to recruit RIPK3 and form a functional necrosome, a central controller of necroptosis^26^. Given that our data also demonstrate DOX-induced phosphorylation of RIPK1 at S161, there can be a critical mechanistic link between ROS accumulation and necroptotic activation in cardiomyocytes. Ser161 autophosphorylation was identified by Yuan et al. in 2008 as a critical positive regulator of RIPK1 kinase activity, and mutation of this site attenuates but does not abolish necroptosis^27^. It is noteworthy that our results showed that DOX-induced RIPK1 activation in MEF cells was associated with phosphorylation of S161 rather than S166, together with the lack of protection by Nec-1 but sensitive to Nec-34 in MEFs. The cooperative inhibition model demonstrated that Nec-1 primarily stabilizes RIPK1 in its inactive DLG-out conformation, while autophosphorylation at S161 represents a distinct activation mechanism: S161 phosphorylation itself stabilizes the active kinase conformation and promotes necrosome assembly, thereby bypassing the conformational state targeted by Nec-1. In contrast, Nec-34 was shown to act through a broader and more cooperative binding mode that can overcome such conformational restrictions^30^, explaining its efficacy in suppressing DOX-induced cell death. The absence of S166 phosphorylation in our MEF system further suggests that DOX triggers a pathway in which RIPK1 activation relies predominantly on S161 autophosphorylation, underscoring S161 as a key regulatory node of necroptosis under oxidative stress conditions. Overall, different cell types responded to DOX by inducing p-S161, p-S166, or both, suggesting that combined treatment with Nec-1 and Nec-34 may provide enhanced efficacy. Moreover, targeting PPP1R3G, which acts upstream of RIPK1 autophosphorylation at S161 and S166, represents an important alternative therapeutic strategy.

Following DOX exposure, mitochondria are the most profoundly affected organelles, with mitochondrial damage occurring as an early event in DOX-induced cardiotoxicity^33^, and limiting DOX-induced mtDNA release mitigates the cardiotoxicity^40^. Interestingly, our results delineate a RIPK1-mtDNA-ZBP1 axis that drives DOX-induced necroptosis. RIPK1 kinase activation promotes mitochondrial DNA (mtDNA) leakage into the cytosol, which elicits type I interferon (IFN-β) production and, in turn, transcriptional upregulation of ZBP1. This notion is supported by public dataset analysis that revealed significant ZBP1 upregulation in DOX-treated cardiomyocytes. Notably, Lei et al. reported that mitochondrial genome instability can drive the accumulation of Z-form mtDNA, which is stabilized by ZBP1 to form a cytosolic complex with cGAS and RIPKs, thereby sustaining, and loss of ZBP1 or IFN-I signaling protects against doxorubicin-induced cardiotoxicity^31^. Collectively, DOX-induced mtDNA instability is not a passive by-product of oxidative damage but an integral, feed-forward driver of necroptotic signaling in cardiomyocytes.

In addition to direct mitochondrial injury, our data indicate that DOX treatment elicits a pronounced inflammatory response characterized by systematically elevated IL-6 and TNFα. Public database mining also showed that doxorubicin exposure upregulates TNF signaling in the heart. Moreover, our findings demonstrate that the presence of TNF, even at very low doses, significantly enhances DOX-induced cell death. These findings are consistent with recent studies showing that adaptive and innate immune components critically contribute to anthracycline cardiotoxicity, and depletion of the pro-inflammatory players alleviates the injury^41,42^. Importantly, as a highly immunogenic form of cell death, necroptosis will lead to the release of DAMPs and inflammatory cytokines, which can further amplify necroptotic signaling and proinflammatory cytokine release (IFN-γ, TNF, CXCL10)^43-45^. In conclusion, necroptosis plays a pivotal role in amplifying DOX-induced cardiac injury by promoting inflammatory cell death and creating a feedforward loop of oxidative stress and inflammation. Targeting necroptosis, especially RIPK1 kinase activity or PPP1R3G/PP1g enzyme complex may offer novel therapeutic opportunities to mitigate the cardiotoxic effects of DOX while preserving its anti-cancer efficacy.

## Conclusion

In summary, our findings establish PPP1R3G as a critical mediator of DOX-induced cardiotoxicity through regulating RIPK1 kinase activation. We propose a model in which, in the presence of PPP1R3G, DOX activates RIPK1, leading to both apoptosis and necroptosis. RIPK1 activation induces IFN-β production, which upregulates ZBP1 and thereby initiates and amplifies late-stage necroptosis. Conversely, PPP1R3G deficiency preserves RIPK1 inhibitory phosphorylation, restrains RIPK1 activation and mtDNA release, attenuates both apoptosis and necroptosis, and protects against cardiac injury in vivo. Together, these results provide mechanistic insight into anthracycline cardiotoxicity and highlight PPP1R3G–RIPK1 signaling as a potential therapeutic target for cardioprotection.

## Acknowledgements

We are grateful to Dr. Dazhi Wang for his valuable guidance and comments, Dr. Ganesh Halade for his technical guidance on mouse echocardiography, and Dr. Chongwu Chi for sharing his expertise in adult mouse cardiomyocyte isolation and culturing. This work is supported by R01GM120502 and R01GM147474.

## Author contributions

Xueling Ma performed most of the experiments, analyzed the data, and drafted the manuscript. Ken Chen helped the mouse work. Zhigao Wang supervised the project, provided critical feedback, and revised the manuscript. All authors reviewed and approved the final version of the manuscript.

