## Supplementary Figures and Tables for "PPP1R3G Deletion Blocks RIPK1-Mediated Apoptosis and Necroptosis in Doxorubicin-Induced Cardiotoxicity"

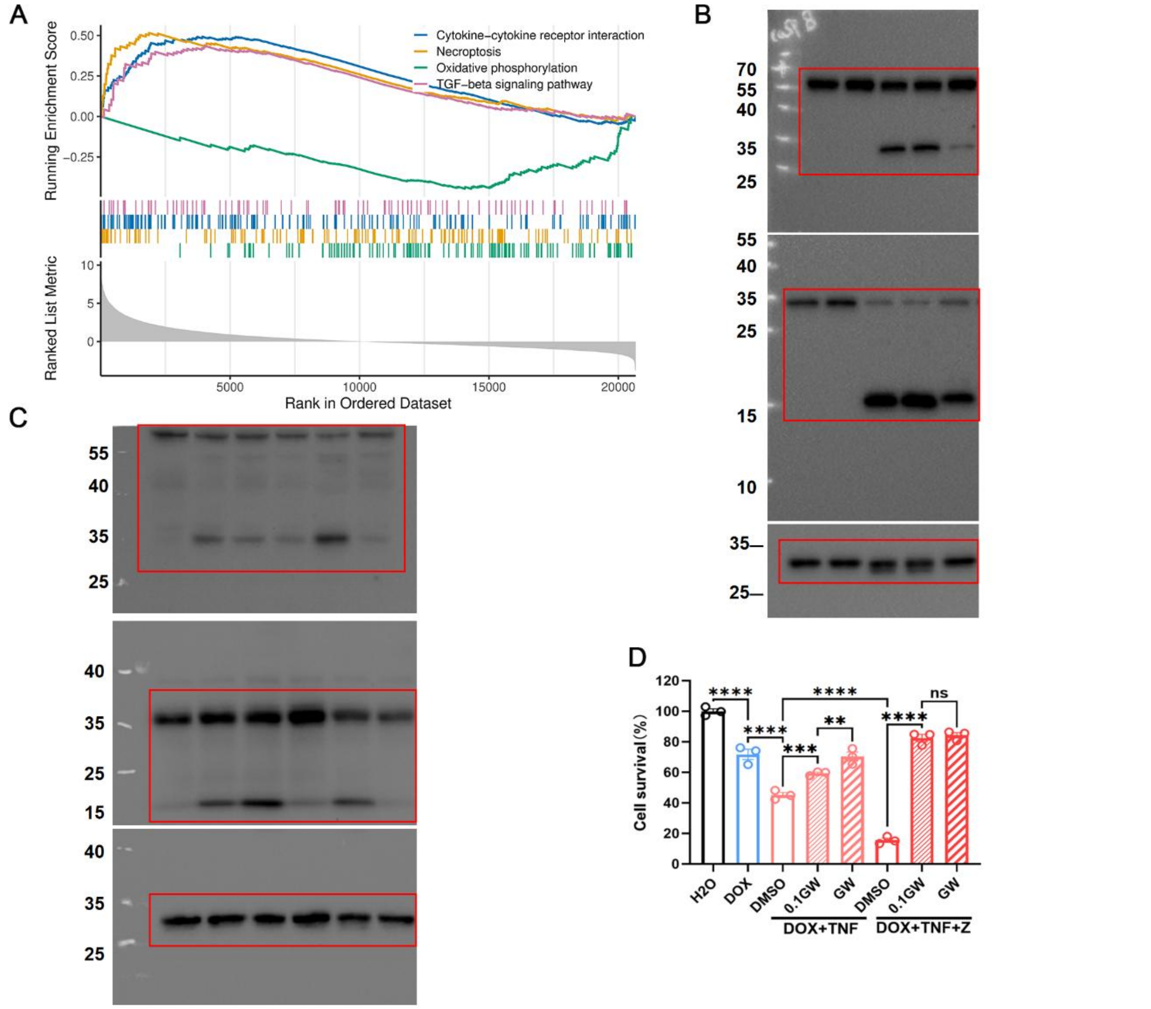


**Figure S1. (A)** GSEA of iPSC-derived cardiomyocytes treated with DOX (GSE217421). **(B, C)** Source data immunoblots of **Fig. 1E, G**. **(D)** Dose-response of GW806742 protection in H9C2 treated with DOX.

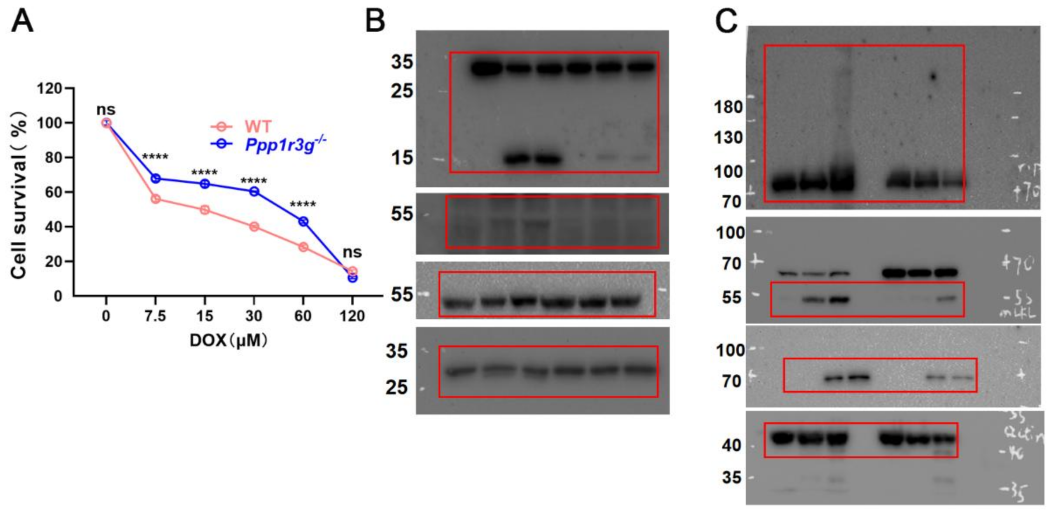


**Figure S2. (A)**Cell survival of WT and *Ppp1r3g*$^{-/-}$ MEFs treated with DOX for 24 hours across different doses. **(B, C)** Source data immunoblots of **Fig. 2E, F**.

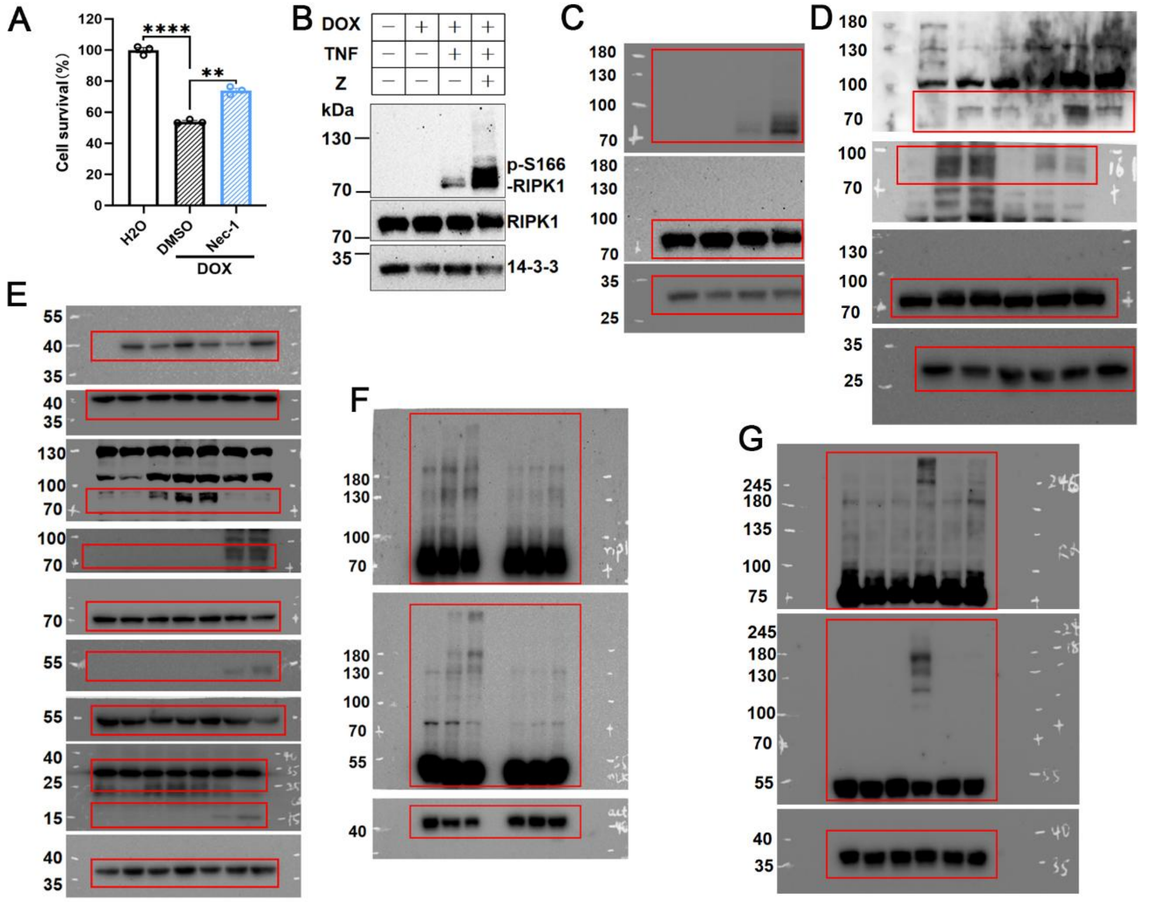


**Figure S3. (A)** Viability of H9C2 cells treated with DOX±Nec-1. **(B)** Immunoblots of RIPK1 phosphorylation after DOX±T±Z in H9C2. **(C-G)** Source data immunoblots of **Fig. S3B, 3A**, **3C-E.**

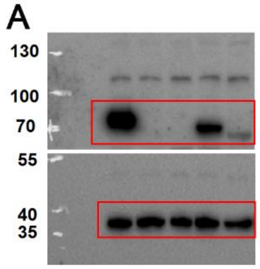


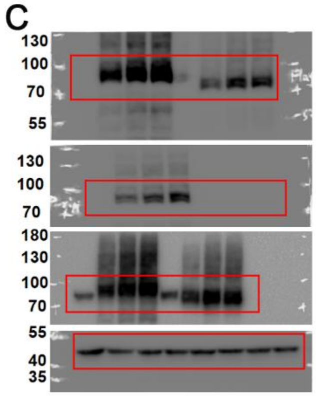


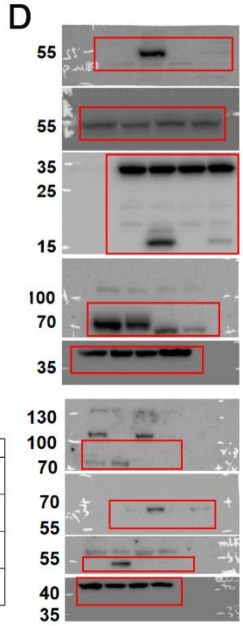


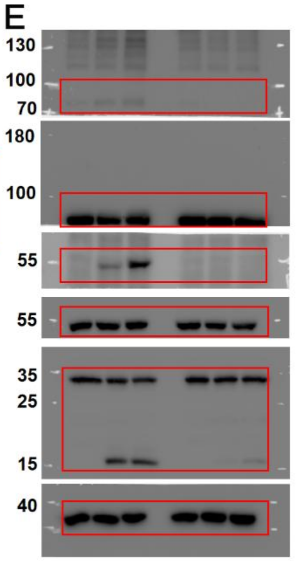


B

| | clone | allele one | allele two |
|---|---|---|---|
| *Ripk1$^{-/-}$* | B-A3 | One nucleotide insertion after nucleotide 158 from ATG | One nucleotide insertion after nucleotide 158 from ATG |
| *Ripk1$^{-/-}$* | D-B6 | One nucleotide insertion after nucleotide 158 from ATG | Five nucleotide deletion after nucleotide 158 from ATG |
| *Ripk1$^{KDD}$* | B-A2 | Two nucleotide deletion after nucleotide 14 from ATG | In frame deletion of aa 6-51 |
| *Ripk1$^{KDD}$* | B-D3 | Ten nucleotide deletion after nucleotide 8 from ATG | In frame deletion of aa 6-53 |

**Figure S4. (A)** Source data immunoblots of **Fig. 4B. (B)** Sequencing results of *Ripk1$^{-/-}$* and *Ripk1$^{KDD}$* clones. **(C-E)** Source data immunoblots of **Fig. 4D, E, G.**

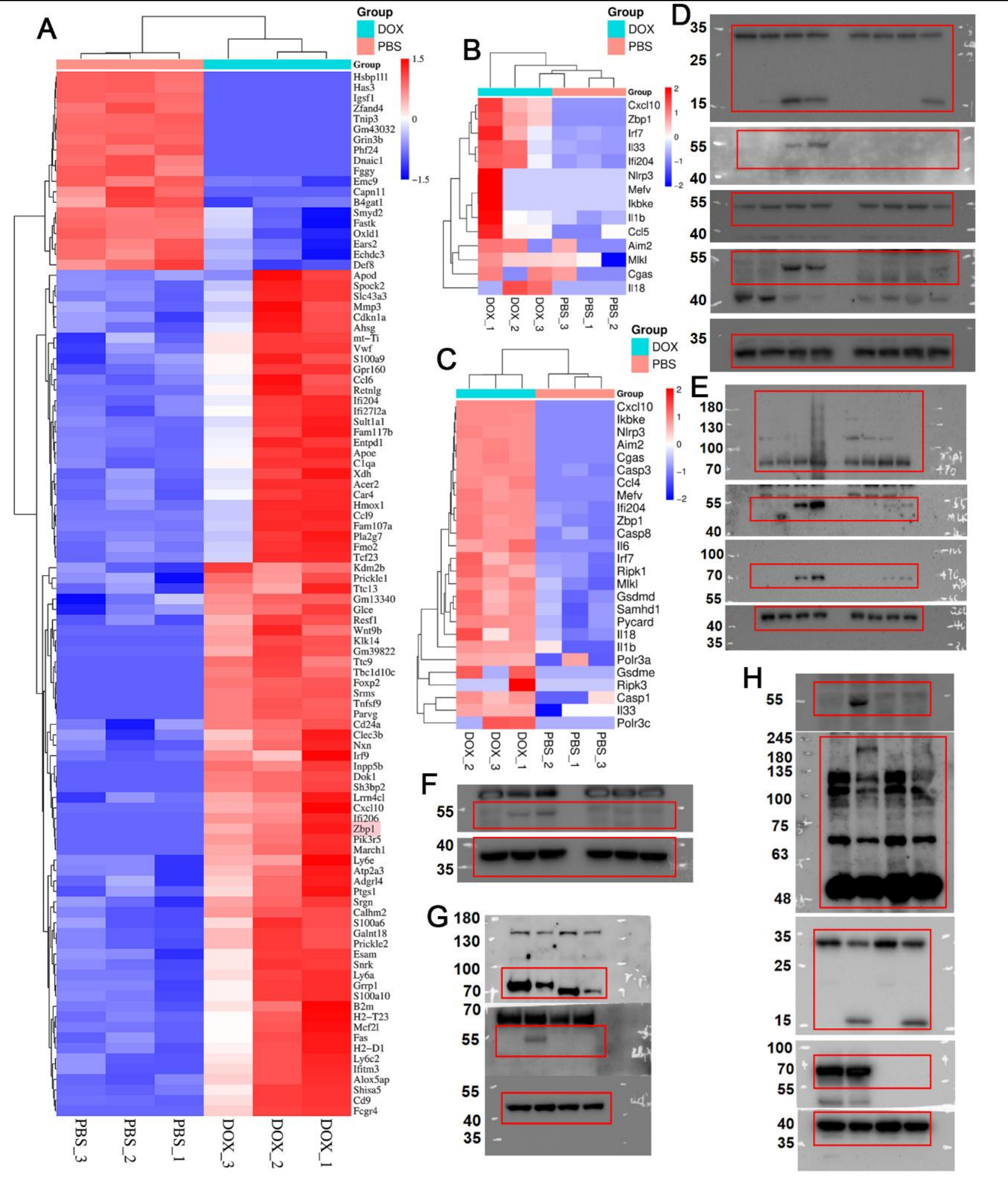


**Figure S5. (A)** Heatmap of RNA-seq results from cardiomyocytes isolated from DOX-treated mice (GSE223698), showing the top 100 ranked genes. **(B, C)** Heatmap of core enrichment genes within the cytosolic DNA-sensing pathway identified from the GSE223698 and GSE226116, respectively. **(D-H)** Source data immunoblots of **Fig. 4C-F, I.**

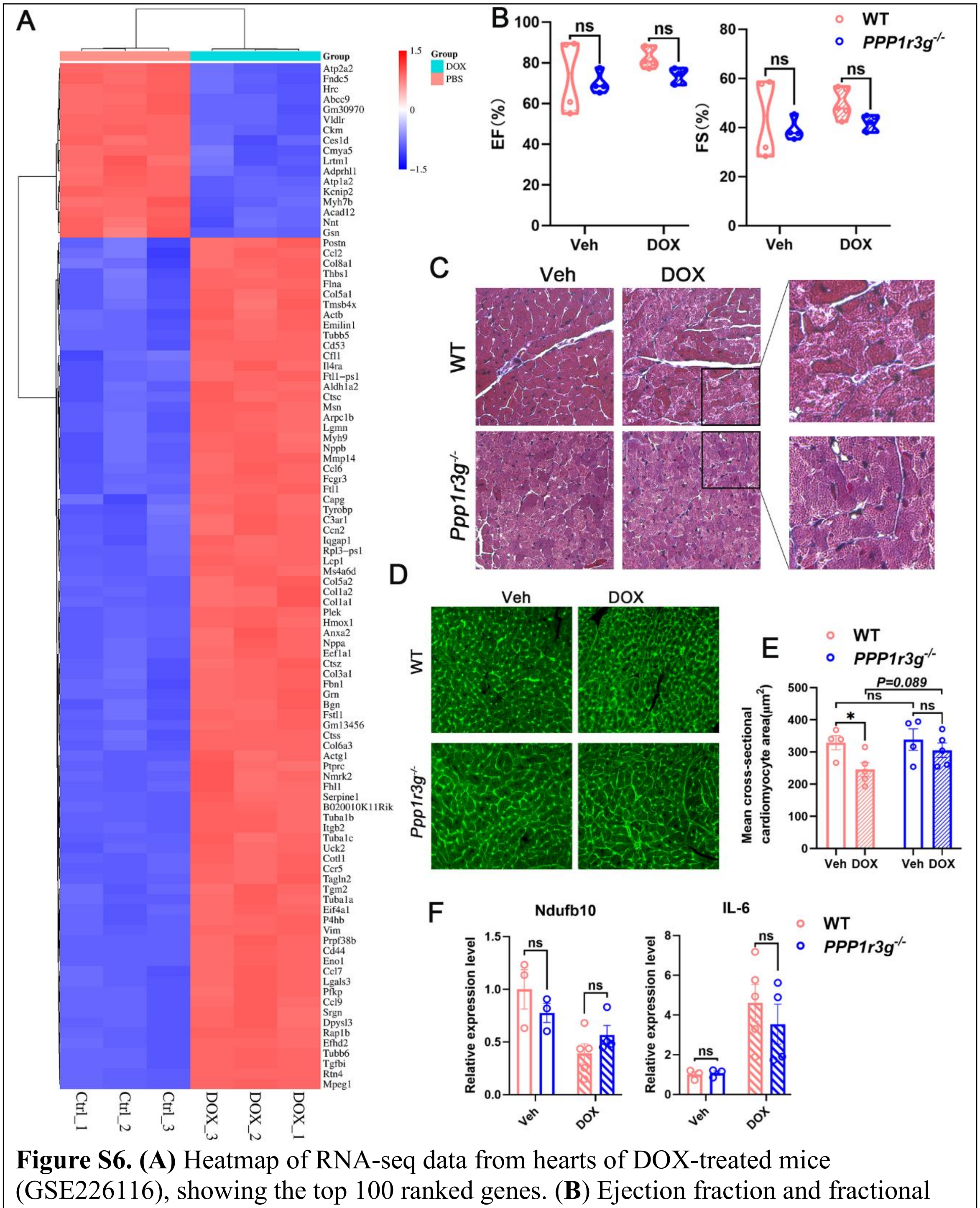


**Figure S6. (A)** Heatmap of RNA-seq data from hearts of DOX-treated mice (GSE226116), showing the top 100 ranked genes. (**B**) Ejection fraction and fractional shortening measured by echocardiography. (**C**) H&E staining of WT and *Ppp1r3g*$^{-/-}$ hearts following DOX treatment, showing marked vacuolization in WT hearts. (**D, E**) WGA staining illustrating changes in cardiomyocyte size. (**F**) RT-qPCR analysis of apical heart tissue from WT and *Ppp1r3g*$^{-/-}$ mice.

**Supplementary Table 1. Core enrichment genes in the cytosolic DNA sensing pathway identified by GSEA (P < 0.05)（GSE226116）**

| Gene symbol | Normalized counts generated by DESeq2 after size factor adjustment | | | | | | log2FoldChange | pvalue |
|---|---|---|---|---|---|---|---|---|
| | DOX_1 | DOX_2 | DOX_3 | PBS_1 | PBS_2 | PBS_3 | | |
| Polr3c | 263.9315466 | 0 | 297.5618628 | 0 | 0 | 0 | 9.856593548 | 0.011646357 |
| Mefv | 102.3408038 | 142.9973573 | 82.05493792 | 0 | 0 | 0 | 9.078399196 | 4.52006E-13 |
| Ikbke | 91.56808759 | 94.72436405 | 100.9906928 | 0 | 0 | 0 | 8.889943042 | 2.82568E-13 |
| Cxcl10 | 97.85217203 | 92.90274166 | 95.58047714 | 0 | 0 | 0 | 8.885205329 | 2.76658E-13 |
| Cgas | 66.43174982 | 72.86489542 | 89.26855884 | 0 | 0 | 0 | 8.560075013 | 7.33103E-12 |
| Aim2 | 59.24993903 | 61.02434992 | 75.74301962 | 0 | 0 | 0 | 8.338466743 | 3.87297E-11 |
| Nlrp3 | 58.35221268 | 71.95408423 | 58.61066995 | 0 | 0 | 0 | 8.285149608 | 5.39473E-11 |
| Il6 | 66.43174982 | 26.41352459 | 32.46129412 | 0 | 0 | 0 | 7.693048307 | 1.65417E-07 |
| Gsdme | 56.55675998 | 57.38110514 | 0 | 0 | 0 | 0 | 7.555657101 | 0.009167758 |
| Ccl4 | 39.49995935 | 32.78920294 | 27.05107844 | 0 | 0 | 0 | 7.357954575 | 1.14107E-07 |
| Casp1 | 126.5794152 | 142.9973573 | 60.41407518 | 0 | 0 | 31.39619078 | 3.395340918 | 0.042385927 |
| Il33 | 161.5907428 | 198.55684 | 155.0928497 | 42.63132089 | 0 | 47.09428616 | 2.522229552 | 0.019760151 |
| Zbp1 | 496.442671 | 565.6137507 | 452.6547125 | 87.44886337 | 112.13784 | 80.733062 | 2.432422045 | 2.44262E-48 |
| Ifi204 | 1081.76025 | 1196.805907 | 1073.026111 | 250.3223714 | 266.3273699 | 214.1668728 | 2.196146147 | 8.2461E-109 |
| Mlkl | 187.6248069 | 187.6271057 | 155.0928497 | 50.28309644 | 71.16439845 | 41.48782353 | 1.700737638 | 1.39589E-11 |
| Casp3 | 184.9316279 | 182.1622386 | 187.5541438 | 64.49353673 | 59.30366537 | 57.18591891 | 1.615390989 | 7.73456E-22 |
| Ripk1 | 241.4883879 | 326.070407 | 272.3141896 | 114.7766332 | 121.8420761 | 95.30986486 | 1.338279632 | 2.86058E-13 |
| Casp8 | 297.1474215 | 325.1595958 | 295.7584576 | 144.2906246 | 125.0768215 | 118.8570079 | 1.241147903 | 9.44747E-22 |
| Samhd1 | 1557.555215 | 1752.400735 | 1389.523729 | 524.6931802 | 803.2951037 | 688.473612 | 1.220575911 | 6.78154E-14 |
| Il18 | 315.9996748 | 416.2407151 | 231.7375719 | 121.3352979 | 162.8155177 | 135.6763959 | 1.198884344 | 2.76715E-06 |
| Irf7 | 2724.599469 | 3200.590531 | 2377.789795 | 1345.619385 | 1363.984304 | 1166.144229 | 1.099003253 | 4.15227E-20 |
| Gsdmd | 238.7952088 | 279.6190362 | 211.9001144 | 101.6593037 | 134.7810577 | 117.7357154 | 1.043587299 | 1.21955E-08 |
| Pycard | 180.4429961 | 191.2703505 | 183.0456308 | 76.51775545 | 101.3553554 | 94.18857233 | 1.027702175 | 4.16326E-10 |

**Supplementary Table 2. Core enrichment genes in the cytosolic DNA sensing pathway identified by GSEA (P < 0.05)（GSE223698）**

| Gene symbol | Normalized counts generated by DESeq2 after size factor adjustment | | | | | | log2FoldChange | pvalue |
|---|---|---|---|---|---|---|---|---|
| | DOX_1 | DOX_2 | DOX_3 | PBS_1 | PBS_2 | PBS_3 | | |
| Mefv | 86.57115801 | 0 | 0 | 0 | 0 | 0 | 20.98642779 | 8.02793E-08 |
| Zbp1 | 655.88657 | 126.7433586 | 54.65038552 | 1 | 1 | 1 | 10.90883152 | 9.96247E-11 |
| Cxcl10 | 196.6192402 | 29.24846736 | 20.7671465 | 1 | 1 | 1 | 9.144515177 | 9.6455E-07 |
| Ifi204 | 1280.959677 | 1156.533147 | 294.0190741 | 156.9713615 | 120.3660872 | 137.7740707 | 2.717628923 | 1.58548E-06 |
| Irf7 | 3288.236697 | 1195.531103 | 607.712287 | 366.8446735 | 337.38979 | 348.7794042 | 2.273212057 | 0.000136528 |
| Il33 | 451.930791 | 370.4805866 | 98.37069394 | 63.30889165 | 60.18304362 | 67.02522358 | 2.271325654 | 0.000232913 |
